# Associations between outdoor temperature and bright sunlight with metabolites in two population-based European cohorts

**DOI:** 10.1101/818740

**Authors:** Boukje C Eveleens Maarse, Nellie Y. Loh, Fredrik Karpe, Frits R Rosendaal, Diana van Heemst, Dennis O Mook-Kanamori, Ko Willems van Dijk, Patrick CN Rensen, Sander Kooijman, Constantinos Christodoulides, Raymond Noordam

## Abstract

**Context:** Outdoor temperature and bright sunlight may directly and/or indirectly modulate systemic metabolism.

**Objective:** We assessed the associations between outdoor temperature and bright sunlight duration with metabolomics.

**Design:** meta-analysis of two cross-sectional studies.

**Setting:** Two population-based European cohort studies.

**Patients or other participants:** Non-diabetic individuals from the Oxford BioBank (OBB; N=6,368; mean age 47.0 years, males 44%) and the Netherlands Epidemiology of Obesity (NEO; N=5,916; mean age 55.6 years, males 43%) studies.

**Intervention(s):** Data on mean outdoor bright sunlight and temperature collected from local weather stations in the week prior to blood sampling.

**Main Outcome Measure(s):** Serum levels of 148 metabolites measured using NMR spectroscopy, including 14 lipoprotein subclasses.

**Statistical analyses:** Multivariable linear regression analyses adjusted for age, sex, body mass index, season and either outdoor temperature or bright sunlight. Summary statistics from the OBB and NEO cohorts were combined using fixed-effect meta-analyses.

**Results:** A higher mean outdoor temperature was associated with increased concentrations of lipoprotein (sub)particles and certain amino acids such as phenylalanine and leucine. In contrast, longer mean hours of bright sunlight were specifically associated with lower concentrations of very low density lipoprotein (sub)particles. The direction of effects was consistent between the OBB and NEO, although effect sizes were generally larger in the OBB.

**Conclusions:** Increased bright sunlight duration is associated with an improved metabolic profile whilst higher outdoor temperature may adversely impact cardiometabolic health.

## Introduction

Outdoor temperature and sunlight intensity and duration affect our daily activities and may consequently have an impact on metabolic health status. Human and animal studies have also highlighted that environmental temperature and sunlight exposure may directly impact systemic metabolism e.g. by modulating brown adipose tissue (BAT) activity and influencing circadian rhythms.[1] Consistent with these notions several epidemiological studies have identified associations between outdoor temperature and prevalence of type 2 diabetes mellitus (T2D) and cardiometabolic diseases.[2–4] For example, Blauw *et al.* showed that the incidence of diabetes was greater in U.S. states with a higher average annual temperature.[4] Similarly, by investigating the associations between outdoor weather conditions and metabolic traits in two European population-based cohorts, namely the Oxford BioBank (OBB) and Netherlands Epidemiology of Obesity (NEO), we have recently shown that increased bright sunlight exposure was associated with a lower Homeostatic Model Assessment for Insulin Resistance (HOMA-IR) and reduced triglyceride levels.[5] In contrast, no associations between mean outdoor temperature and glucose or lipid metabolism were detected. These results indicate that outdoor sunlight is specifically associated with a more beneficial cardiometabolic risk profile, although further studies are necessary to confirm these findings and determine the underlying biological mechanism(s) underpinning these associations.

Recently, high-throughput metabolite profiling has emerged as a powerful tool for the exploration of disease mechanisms and the identification of novel therapeutic targets[6–9], especially in the context of (cardio)metabolic disorders.[10, 11] Based on our earlier study[5], we hypothesized that outdoor bright sunlight would be associated with a more favourable metabolite profile. In this study, we performed a cross-sectional analysis investigating the association between outdoor bright sunlight and environmental temperature with plasma levels of 148 lipids and metabolites as determined by NMR spectroscopy in a combined sample of over 12,000 middle-aged population-based subjects, without pre-existing diabetes mellitus, from the OBB and NEO study cohorts.

## Materials and Methods

### Study design

The OBB is a population-based cohort of randomly selected healthy participants aged 30 to 50 years from Oxfordshire (UK). Individuals with a history of myocardial infarction, diabetes mellitus, heart failure, untreated malignancy, other ongoing systemic diseases or ongoing pregnancy were not eligible for study inclusion. Participants were included between 1999 and May 2015. The OBB cohort comprises 7,185 individuals (4,054 women and 3,131 men). A more detailed description of the study recruitment criteria and population characteristics is reported elsewhere.[12]

The NEO study is population-based prospective cohort study of men and women aged between 45 and 65 years with an oversampling of individuals with a BMI of 27 kg/m^2^ or higher, living in the greater area of Leiden (in the West of the Netherlands). In addition, all inhabitants aged between 45 and 65 years from one municipality (Leiderdorp) were invited in the study irrespective of their BMI, to allow for a reference distribution of BMI. Between September 2008 and September 2012, 6,671 individuals (3,505 women and 3,168 men) were included in the study. Detailed information about the study design and data collection has been described previously[13].

In both cohorts, participants were invited for a detailed baseline assessment, conducted after an overnight fast, which included blood sampling and anthropometry. Both studies were approved by local ethics committees, and written informed consent was obtained from all study participants.

### Study population

In the OBB, we excluded individuals with missing data on mean outdoor temperature and/or bright sunlight in the week preceding the centre visit, body composition, and fasting metabolomics (missing in 817 individuals). Consequently, data from 6,368 individuals were used for the present analyses. From NEO, we excluded individuals with both treated and diagnosed diabetes, as well as subjects with a fasting glucose concentration above 7.0 mmol/L (N = 749) in order to have a uniform population as that of the OBB regarding glycaemic status. Additionally, we excluded participants who were nonfasting, or had missing data on mean outdoor temperature and/or bright sunlight in the week preceding the centre visit, body composition, and fasting metabolomics (missing in 6 individuals). As a result, a total of 5,916 individuals were used for the analyses presented in this study.

### Data collection on outdoor temperature and bright sunlight

Data on the mean temperature and hours of bright sunlight (defined as global radiation >120 W/m^2^) were collected from the weather station that was located closest located to either Oxfordshire or Leiden. Based on these data, we estimated the mean outdoor temperature and bright light over the week prior to the date of the blood sampling. For the OBB data were obtained from the Radcliffe Meteorological Station (Woodstock Road, Oxford, UK). For the NEO study we obtained data from a measurement station from the Koninklijk Nederlands Meteorologisch Instituut (Royal Dutch Meteorological Institute).

### NMR-based metabolic biomarker profiling

We used a high-throughput proton NMR metabolomics platform[14] (Nightingale Health Ltd., Helsinki, Finland) to quantify 148 lipid and metabolite concentrations in fasting serum samples. The NMR spectroscopy was conducted at the Medical Research Council Integrative Epidemiology Unit (MRC IEU) at the University of Bristol, Bristol, United Kingdom, and processed by Nightingale’s biomarker quantification algorithms (version 2014). This method provides quantification of lipoprotein subclass profiling with lipid concentrations within 14 lipoprotein subclasses. The 14 subclass sizes were defined as follows: extremely large VLDL with particle diameters from 75 nm upwards and a possible contribution of chylomicrons, five VLDL subclasses (average particle diameters of 64.0 nm, 53.6 nm, 44.5 nm, 36.8 nm, and 31.3 nm), IDL (28.6 nm), three LDL subclasses (25.5 nm, 23.0 nm, and 18.7 nm), and four HDL subclasses (14.3 nm, 12.1 nm, 10.9 nm, and 8.7 nm). Within the lipoprotein subclasses the following components were quantified: total cholesterol, total lipids, phospholipids, free cholesterol, cholesteryl esters, and triglycerides. The mean size for VLDL, LDL and HDL particles was calculated by weighting the corresponding subclass diameters with their particle concentrations. Furthermore, 47 metabolic measures were determined that belong to classes of apolipoproteins, cholesterol, fatty acids, glycerides, phospholipids, amino acids, fluid balance, glycolysis-related metabolites, inflammation, and ketone bodies. Detailed experimentation and applications of the NMR metabolomics platform have been described previously,[14] as well as representative coefficients of variations (CVs) for the metabolic biomarkers.[15] Full names and descriptions of metabolic measures are listed in Supplementary Table 1.

### Covariates

Height and weight were measured by research nurses at the OBB and NEO study centres. BMI was calculated by dividing the weight in kilograms by the height in meters squared. Season was derived from the date of the blood sampling (winter: December – February, spring: March – May, summer: June – August, autumn: September – November). In both cohorts, use of lipid-lowering medication was determined by medication inventory.

### Statistical analysis

In the NEO study, participants with a BMI of 27 kg/m^2^ or higher are oversampled. To correctly represent associations for the general population[16], we corrected for oversampling of participants with a BMI ≥ 27 kg/m^2^, which was done by weighting individuals towards the BMI distribution of participants from the Leiderdorp municipality[17], whose BMI distribution was similar to the BMI distribution of the general Dutch population[18]. Consequently, all results were based on weighted analyses and results apply to a population-based study without oversampling of individuals with a BMI ≥ 27 kg/m^2^.

All analyses were performed using STATA version 12.1 (StataCorp LP, TX, US). Baseline characteristics of the OBB and NEO study populations were conducted separately. Continuous variables were expressed as (weighted) mean (with standard deviation [SD] for normally distributed variables or (weighted) median (inter quartile range [IQR]) for skewed variables. Dichotomous or categorical variables were expressed as (weighted) proportion (%).

As most of the metabolic outcome variables were not normally distributed, we log-transformed all these variables prior to standardization to a standard normal distribution (mean = 0, s.d. = 1) to be able to better compare the effect sizes of the different study outcomes. Associations of mean bright sunlight and temperature with lipid and metabolite concentrations were examined using multivariable linear regression analyses for the combined population of the OBB and NEO study populations. Estimates retrieved from the analyses were subsequently meta-analysed using fixed-effects meta-analysis as implemented in the rmeta statistical package in R. For presentation purposes, we analysed the data per 5 degrees Celsius increase in outdoor temperature and per hour increase in bright sunlight exposure. Consequently, results can be interpreted as the difference in standard deviation per unit increase in either outdoor temperature (5 degrees Celsius) and bright sunlight exposure (1 hour).

To study the impact of adjustment of several of the covariates in the multivariable linear regression analyses, we considered 3 different statistical models. Model 1 was adjusted for age, sex and BMI (given our earlier observation that the weather exposures were associated with BMI).[5] Model 2 was additionally adjusted for season. Model 3 was additionally adjusted either for the mean temperature or mean hours of bright sunlight to fully dissect the two weather exposures in the present study, which were moderately correlated with each other.[5] In a sensitivity analysis, we additionally excluded individuals who used cholesterol-lowering treatment. To test the consistency of the results between the OBB and the NEO cohorts, a plot was constructed with beta estimates of the identified metabolites in both cohorts visualized against each other for outdoor temperature and bright sunlight, using the R-package ggplot2.[19]

Given the high number of statistical tests performed in the present study, we corrected for multiple testing. The metabolic biomarkers used for the present study are correlated with each other, and therefore, conventional correction for multiple testing (e.g., Bonferroni) is too stringent. To obtain the number of independent metabolic biomarkers, we used the method as described by Li et al.,[20] which takes the correlation between the different metabolic biomarkers into account. Based on this method, we found 37 independent metabolic markers. For this reason, associations were considered to be statistically significant in case the *p*.-value was below 0.00134 (i.e. 0.05/37).

## Results

### Study population characteristics

The total study population (N = 12,284) comprised of 6,368 individuals from the OBB and 5,916 individuals from the NEO study (see Table 1). Compared to participants from the NEO cohort, OBB volunteers were younger (mean age 47.0 vs. 55.6 years) and had a lower mean BMI (25.9 vs 26.1 kg/m^2^). Mean outdoor temperature during the week prior to blood sampling was similar between the two cohorts (10.7 degrees Celsius in both cohorts) although mean hours of bright sunlight were higher in Leiden than in Oxfordshire (3.8 vs 5.0 hours).

**Table 1:** Characteristics of the study populations

|  | OBB (N = 6,368) | NEO (N = 5,916) |
| --- | --- | --- |
| Age in years, mean (SD) | 47.0 (7.0) | 55.6 (6.0) |
| Men, % | 43.8 | 43.3 |
| Body mass index in kg/m<sup>2</sup>, mean (SD) | 25.9 (4.6) | 26.1 (4.3) |
| Winter, % | 24.1 | 23.9 |
| Spring, % | 26.3 | 26.3 |
| Summer, % | 25.2 | 24.3 |
| Autumn, % | 24.5 | 25.5 |
| Outdoor temperature (7 days) in °C, mean (SD) | 10.7 (6.8, 15.1) | 10.7 (6.9, 15.5) |
| Outdoor bright sunlight (7 days) in hours, median (IQR) | 3.8 (2.3, 5.8) | 5.0 (2.7, 7.0) |

Results of the NEO study population are weighted towards the body mass distribution of the general population. Abbreviations: IQR, interquartile range; NEO, Netherlands Epidemiology of Obesity; N, number of participants; SD, standard deviation.

### Outdoor temperature and serum metabolites

A high-throughput proton NMR metabolomics platform[14] was utilized to quantify 148 lipid and metabolite concentrations in serum samples obtained following an overnight fast. Associations between outdoor temperature and circulating levels of these metabolites following adjustment for age, sex, BMI, season and hours of bright sunlight in the combined study population are presented in **Figure 1**. In total we identified 27 metabolites whose concentration was associated with outdoor temperature after correction for multiple testing (*p*.< 1.34e^-3^). For example, a higher mean outdoor temperature in the week preceding blood sampling was associated with an increased concentration of total cholesterol (β (SE) = 0.0642 (0.0184) SD per 5 degrees Celsius, *p*.=5.03e^-4^). Lipoprotein subfraction analysis revealed that higher outdoor temperature was most strongly associated with raised serum very small VLDL cholesterol (XS-VLDL; 0.0605 (0.0185) per 5 degrees Celsius; *p*.=1.00e^-3^) and IDL cholesterol (0.0604 (0.0185) SD per 5 degrees Celsius, *p*.=1.12e^-3^) particles and their subcomponents. No associations between environmental temperature and serum LDL cholesterol (sub)particles were detected although, a higher mean outdoor temperature was positively associated with increased medium HDL cholesterol (M-HDL) particle concentration (0.0629 (0.0175) SD per 5 degrees Celsius, *p*.=3.34e^-3^).

**Figure 1:**
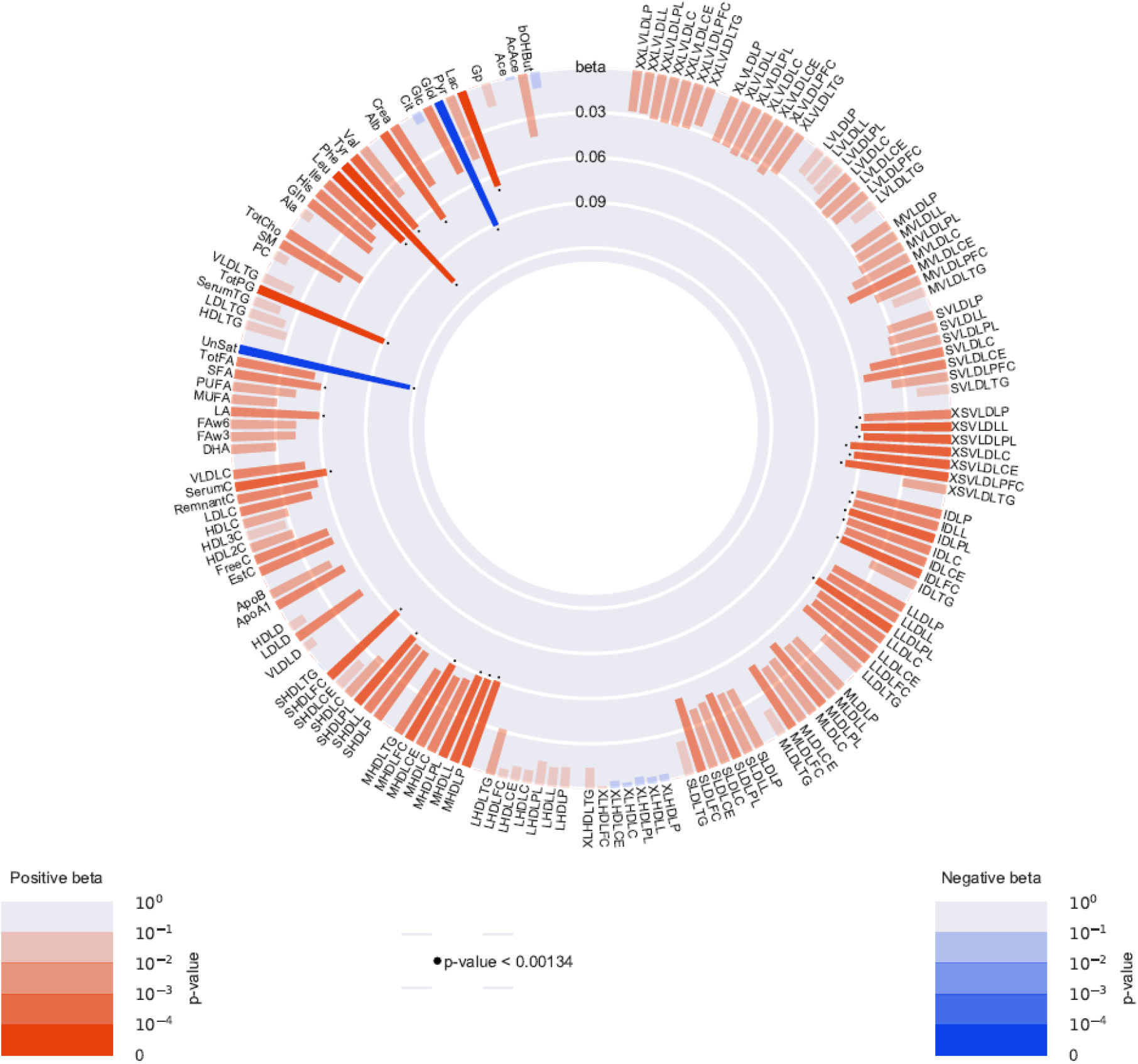
Observational associations between outdoor temperature and 148 lipid and metabolite biomarkers, in a meta-analysis of the Oxford Bio Bank (OBB) study (N = 6,368) and Netherlands Epidemiology of Obesity (NEO) study (N = 5,916). Bar heights represent the magnitude of the beta-coefficient from linear regression, which is expressed as mean SD difference in metabolic measure per 5°C. Red bars indicate positive betas and blue bars indicate negative betas. The transparency of the bars indicates the level of statistical significance. A *p*.-value <1.34e^-3^ is considered as statistical significant, as represented by the black dots. Results are adjusted for age, sex, BMI, season and mean hours of bright sunlight. Full names and descriptions of metabolic measures are listed in Supplementary Table 1. Results of the NEO study population are weighted towards the body mass distribution of the general population.

In addition to increased serum lipid particles, a higher mean outdoor temperature was associated with lower circulating concentrations of unsaturated fatty acids (−0.1212 (0.0186) SD per 5 degrees Celsius, *p*.=7.20e^-11^). Finally, positive associations were detected between mean outdoor temperature and serum concentrations of the amino acids leucine (0.0660 (0.0165) SD per 5 degrees Celsius, *p*.=6.44e^-05^), phenylalanine (0.1108 (0.0182) SD per 5 degrees Celsius, *p*.=1.25e^-09^) and tyrosine (0.0657 (0.0178) SD per 5 degrees Celsius, *p*.=2.23e^-4^) as well as, the anaerobic glycolysis end-product lactate (−0.0702 (0.0177) SD per 5 degrees Celsius, *p*.=7.64e^-5^). Directionally similar associations were detected in the OBB and NEO cohorts (**Figure 2**) although, effect sizes were generally stronger in the OBB. Results did not materially differ after excluding individuals using cholesterol-lowering medications (results not shown).

**Figure 2:**
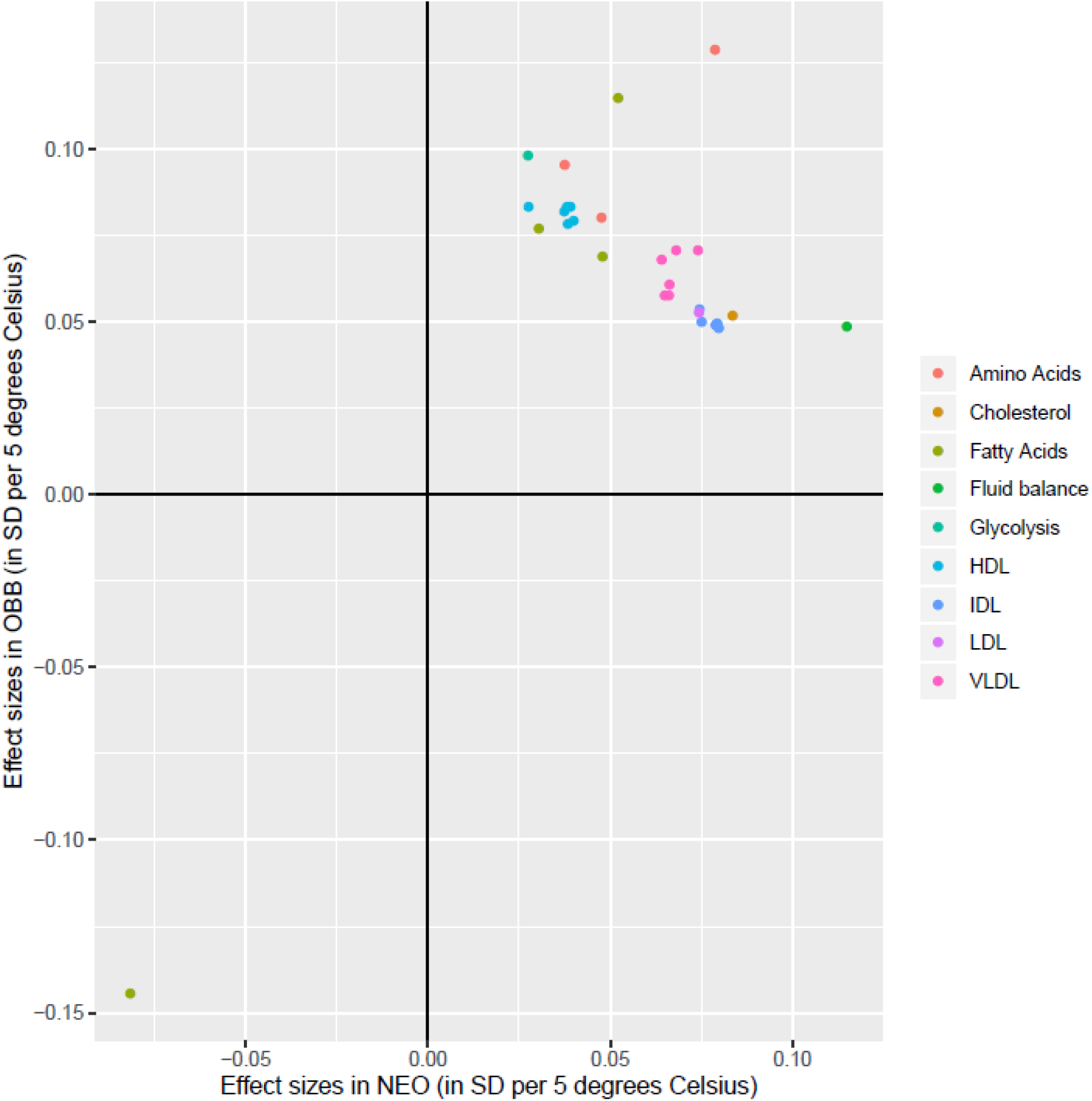
Correlations between the SD of the effect sizes per 5 degrees Celsius of the OBB and the NEO study. Only metabolites that were significantly associated (*p*.<1.34e^-3^) with outdoor temperature were included. X-axis presents the beta estimates per 5 degrees Celsius in the Netherlands Epidemiology of Obesity (NEO) study; Y-axis presents the beta estimates per 5 degrees Celsius in Oxford Biobank (OBB).

### Bright sunlight and serum metabolites

Associations between outdoor bright sunlight and NMR spectroscopy measured metabolites following adjustment for age, sex, BMI, season and outdoor temperature in the combined OBB and NEO population are presented in **Figure 3**. After accounting for multiple testing (*p*.< 1.34e^-3^) mean outdoor bright sunlight was significantly associated with a total of 58 metabolites. In contrast to the pattern of associations detected with increased temperature, a longer duration of bright sunlight during the week preceding the study visit was associated with decreased serum levels of total cholesterol (−0.0194 (0.0053) SD per 1 hour bright sunlight, *p*.=2.36e^-4^), VLDL cholesterol (−0.0243 (0.0054) SD per 1 hour bright sunlight, *p*.=8.06e^-6^) and remnant cholesterol (−0.0241 (0.0054) SD per 1 hour bright sunlight, *p*.=8.43e^-6^). More specifically, we found lower serum concentrations of extremely large VLDL cholesterol (XXL-VLDL: −0.0203 (0.0049) SD per 1 hour bright sunlight, *p*.=3.28e^-5^), very large VLDL cholesterol (XL-VLDL: −0.0199 (0.0049) SD per 1 hour bright sunlight, *p*.=5.76e^-5^), very small VLDL cholesterol (XS-VLDL: −0.0222 (0.0055) SD per 1 hour bright sunlight, *p*.=5.26e^-5^) and IDL cholesterol (−0.0189 (0.0054) SD per 1 hour bright sunlight, *p*.=4.53e^-4^) particles and their subcomponents. No association between bright sunlight duration and serum LDL cholesterol concentration was detected. The reduction in VLDL and IDL cholesterol particle numbers was also reflected in the negative association between hours of bright sunlight and serum levels of ApoB (−0.0201 (0.0054) SD per 1 hour bright sunlight, *p*.=2.14e^-4^).

**Figure 3:**
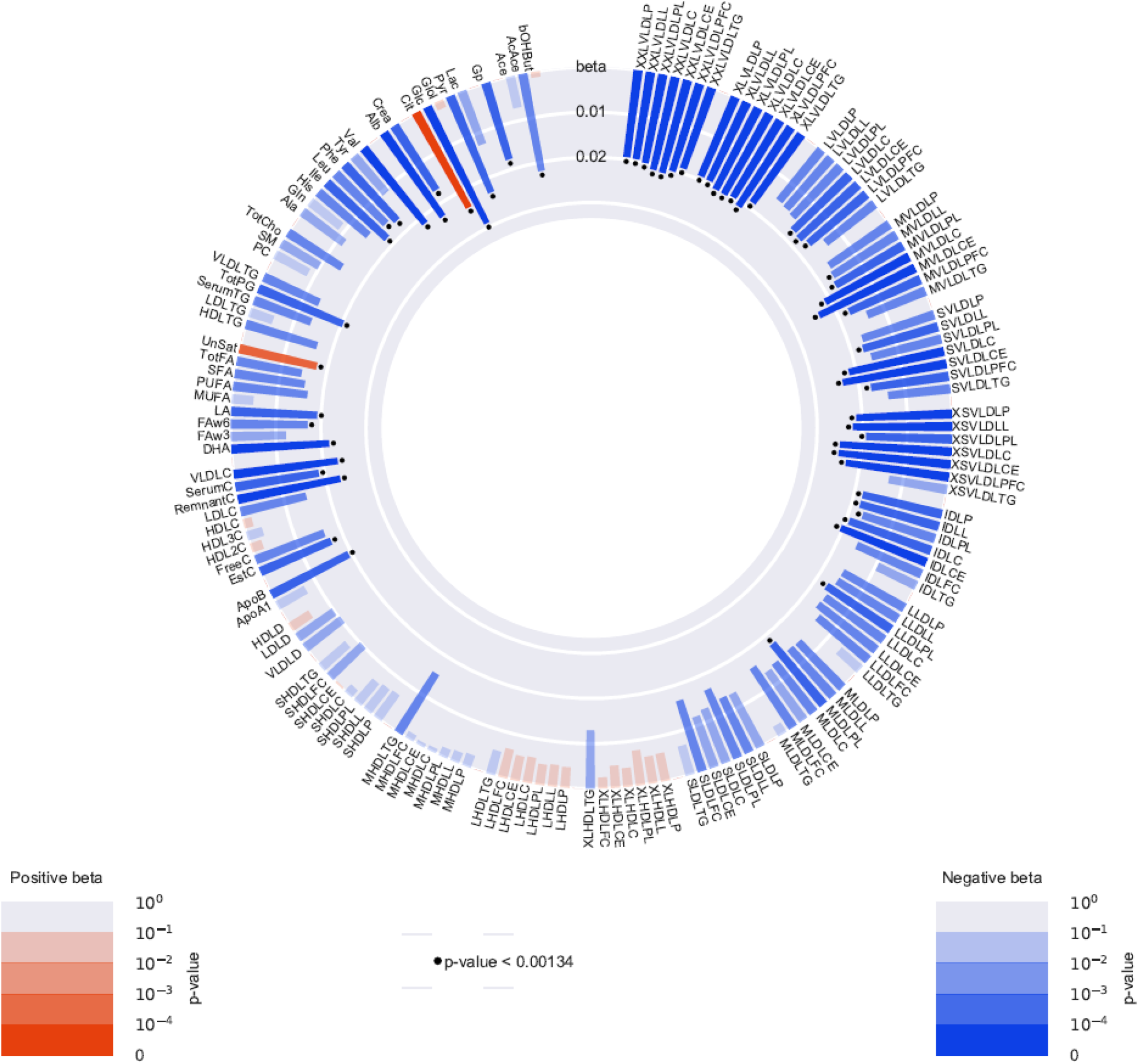
Observational associations between bright sunlight and 148 lipid and metabolite biomarkers, in a meta-analysis of the Oxford Bio Bank (OBB) study (N = 6,368) and Netherlands Epidemiology of Obesity (NEO) study (N = 5,916). Bar heights represent the magnitude of the beta-coefficient from linear regression, which is expressed as mean SD difference in metabolic measure per hour of sunlight. Red bars indicate positive betas and blue bars indicate negative betas. The transparency of the bars indicates the level of statistical significance. A *p*.-value <1.34e^-3^ is considered as statistical significant, as represented by the black dots. Results are adjusted for age, sex, BMI, season and mean outdoor temperature. Full names and descriptions of metabolic measures are listed in Supplementary Table 1. Results of the NEO study population are weighted towards the body mass distribution of the general population.

In addition to lower serum VLDL and IDL lipoprotein particle concentrations, longer bright sunlight duration was associated with an increased systemic concentration of unsaturated fatty acids (0.0183 (0.0055) SD per 1 hour bright sunlight, *p*.=8.19e^-4^). Finally, negative associations between ambient bright sunlight hours and serum levels of the amino acid phenylalanine (−0.0176 (0.0053) SD per 1 hour bright sunlight, *p*.=9.12e^-4^), and the branched-chain amino acids isoleucine (−0.0189 (0.0052) SD per 1 hour bright sunlight, *p*.=2.72e^-4^), leucine (−0.0164 (0.0050) SD per 1 hour bright sunlight, *p*.=1.15e^-3^ 0.0012) and valine (−0.0224 (0.0051) SD per 1 hour bright sunlight, *p*.=1.15e^-5^) were also detected. Indeed, bright sunlight duration was generally inversely associated with the circulating levels of all the amino acids quantified by the NMR metabolomics platform utilised although these associations were not significant after correction for multiple testing.

As shown in **Figure 4**, with the notable exception of leucine, glucose, glycoprotein, acetyl and total phosphoglyceride levels the effects of bright sunlight on the plasma metabolome were directionally identical in the OBB and NEO datasets.

**Figure 4:**
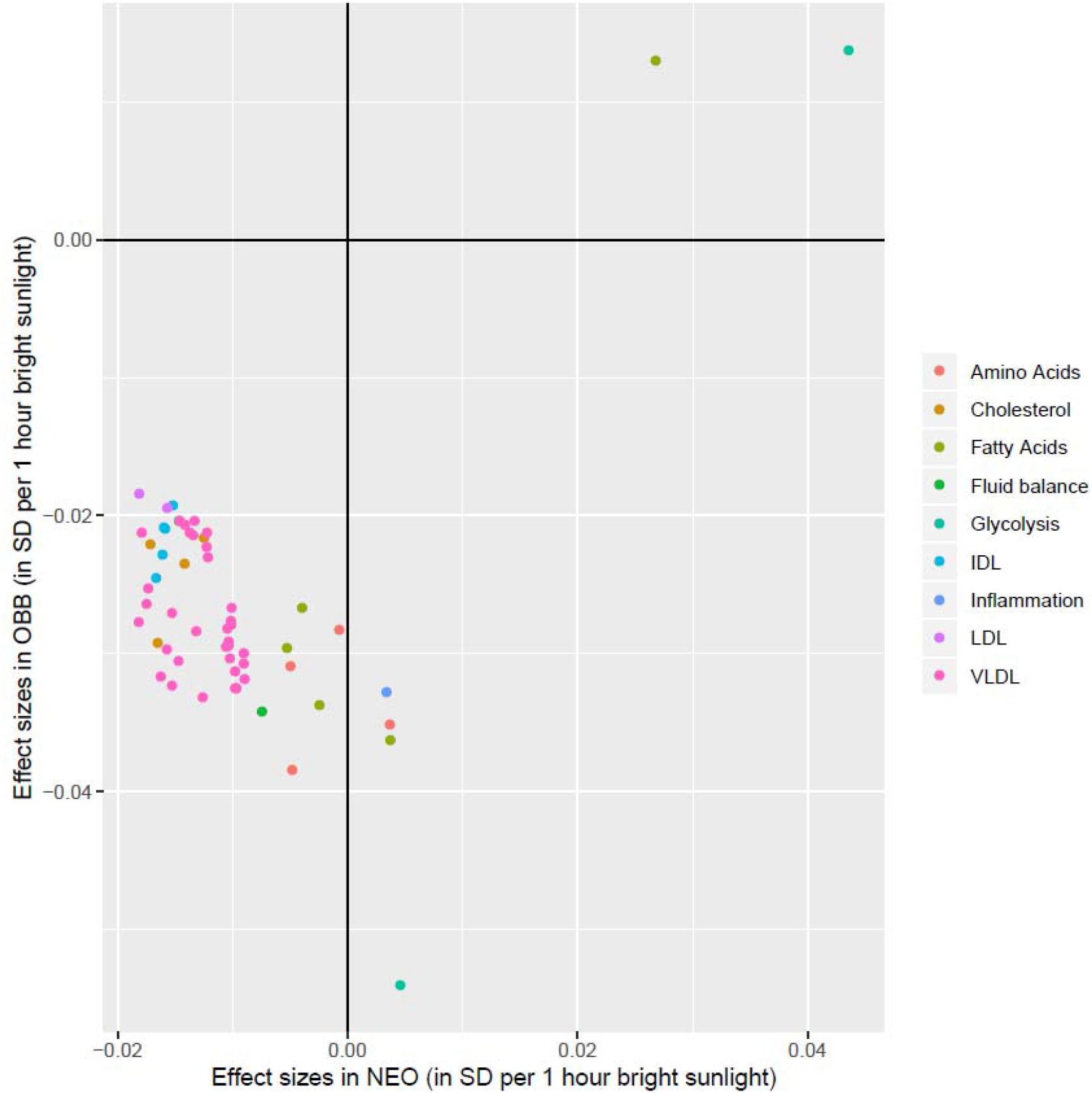
Correlations between the SD of the effect sizes per hour of bright sunlight of the OBB and the NEO study. Only metabolites that were significantly associated (*p*.<1.34e^-3^) with bright sunlight were included. X-axis presents the beta estimates per hour bright sunlight in the Netherlands Epidemiology of Obesity (NEO) study; Y-axis presents the beta estimates per hour bright sunlight in Oxford Biobank (OBB).

## Discussion

We compiled data from two population-based European cohorts, comprising a combined sample size of more than 12,000 non-diabetic individuals, to investigate the associations between mean outdoor temperature and bright sunlight and serum metabolites. Our results suggest that whilst higher outdoor temperature is associated with an unfavourable metabolic profile[21–23], prolonged exposure to bright sunlight is in contrast characterized by lower plasma lipoprotein concentrations and potentially decreased circulating levels of BCAAs.

Our findings build on our previous study in which we reported that bright sunlight, but not outdoor temperature was associated with enhanced glucose and lipid metabolism, as determined by a lower HOMA-IR and decreased plasma triglycerides.[5] Extending these findings we now show that prolonged mean hours of bright sunlight are correlated with lower levels of serum cholesterol, ApoB and the branched-chain amino acids (BCAAs) isoleucine, leucine and valine. The latter result has to be taken with caution as the effect sizes of the associations between bright sunlight and BCAAs differed considerably between the OBB and the NEO cohorts, and had opposite directions in case of leucine. Furthermore, we demonstrate that the association between bright sunshine and lower serum cholesterol concentration was primarily driven by a reduction in VLDL particle numbers, reflecting either decreased VLDL production or increased VLDL clearance.[24] Consistent with our earlier study[5] a negative association was also detected between bright sunlight and plasma triglycerides (p-value<0.05) although this did not survive correction for multiple testing. Notably, with a few exceptions, associations between outdoor temperature and circulating metabolites were directionally opposite to those detected with bright sunlight, suggesting that increased environmental temperature is associated with an unfavourable cardiometabolic profile. In this respect, higher circulating concentrations of BCAAs and aromatic amino acids, total VLDL, total cholesterol and ApoB have been associated with an increased risk of incident T2D.[21, 23] Additionally, systemic BCAAs have been shown to be causally related to the development of T2D[25] whilst increased plasma levels of VLDL cholesterol are known to be associated with poor glycaemic control in patients with T2D.[23] As well as (recently) being linked to the development of T2D, increased plasma levels of VLDL, IDL and LDL lipoprotein particles, in addition to cholesterol remnants and ApoB, are established risk factors for incident myocardial infarction and ischemic stroke.[22] Therefore, the results of our study suggest that during periods of increased bright sunlight, individuals experience a lower risk of cardiometabolic diseases which maybe predominantly attributable to lower concentrations of VLDL (sub)particles. In contrast, the reverse might be true during spells of high environmental temperature.

We previously raised the hypothesis that the association between bright sunlight and a favourable metabolic profile is mediated, at least in part, by the influence of sunlight on melatonin metabolism.[5] Consistent with this notion nocturnal melatonin concentrations increase with prolonged exposure to bright sunlight[26] and melatonin has been shown to play an important role in the circadian rhythm of insulin secretion by pancreatic β-cells.[27] Moreover, common and rare genetic variants at *MTNR1B,*. the melatonin receptor gene are strongly associated with fasting glucose levels and T2D-risk[28, 29] whilst treatment with melatonin was associated with an improved lipid profile and blood pressure in patients with the metabolic syndrome and post-menopausal women.[30] PMID: 18298467. To further investigate this hypothesis, future studies investigating the effects of light therapy on melatonin level and systemic metabolism are necessary.[31] Another potential mechanism for the observed associations between bright sunlight and a favourable cardiometabolic profile could involve vitamin D metabolism.[32] However, Mendelian Randomization studies have shown a lack of a causal association between circulating vitamin D levels and T2D as well as lipid levels and cause-specific vascular disease and mortality[33, 34] Nevertheless, these are merely hypotheses that should be further explored in future studies incorporating larger sample sizes.

A possible mechanism underlying the associations between increased outdoor temperature and an unfavourable metabolite profile is impaired BAT activity. Brown adipocytes are activated by cold and combust intracellularly stored triglycerides to generate heat.[35] Additionally, it has been shown that activated BAT takes up glucose and lipids to supplement its intracellular triglyceride stores[36, 37] and that both the presence and activity of BAT are negatively associated with outdoor temperature.[38, 39] A strong association was also found between winter season and the presence and activity of BAT.[39, 40] Based on these data, it is possible that a higher outdoor temperature leads to an unfavourable glucose and lipid profile by means of diminished BAT. Nevertheless, testing of this hypothesis is complicated given the likely dilution effect of clothing and indoor heating on BAT activation during spells of cold environmental temperature in real-life.

A particular strength of our study is the combination of two large independent cohorts originating from different countries and comprising more than 12,000 participants to study the effects of temperature and bright sunlight on metabolism. A limitation is the inherent heterogeneity in the two cohorts, which has to be taken into account in interpreting the results. These differences might have been driven by differences in lifestyle between the United Kingdom and The Netherlands. Unfortunately, detailed data on lifestyle such as habitual food intake and physical activity were not available in both cohorts. Consequently, we are also not able to investigate whether these lifestyle factors potentially mediated the association between either outdoor temperature and/or bright sunlight and metabolite concentrations. Nevertheless, additional adjustment for season, which would partially adjust for variations in lifestyle factors, did not materially alter our findings.

In summary, extending our earlier work[5] we provide further evidence that increased bright sunlight is associated with improved cardiometabolic health. In contrast increased outdoor temperature may be associated with an adverse cardiometabolic profile. Future research is required to elucidate the mechanistic basis of the observed associations.

## Acknowledgements and funding

We express our gratitude to all individuals who participate in the OBB and NEO study. We are grateful to all participating general practitioners for inviting eligible participants and all research nurses for collecting the data. The Oxford Biobank is supported by the NIHR Oxford BRC Obesity and Lifestyle theme. The NEO study is supported by the participating Departments, the Division and the Board of Directors of the Leiden University Medical Center, and by the Leiden University, Research Profile Area ‘Vascular and Regenerative Medicine’. FK is supported by British Heart Foundation (RG/17/1/32663). CC is a BHF Intermediate clinical research fellow. SK is supported by the Dutch Heart Foundation (2017T016). We acknowledge the support from the Netherlands Cardiovascular Research Initiative: an initiative with support of the Dutch Heart Foundation (CVON2014-02 ENERGISE).

---

**Supplementary Table 1.**
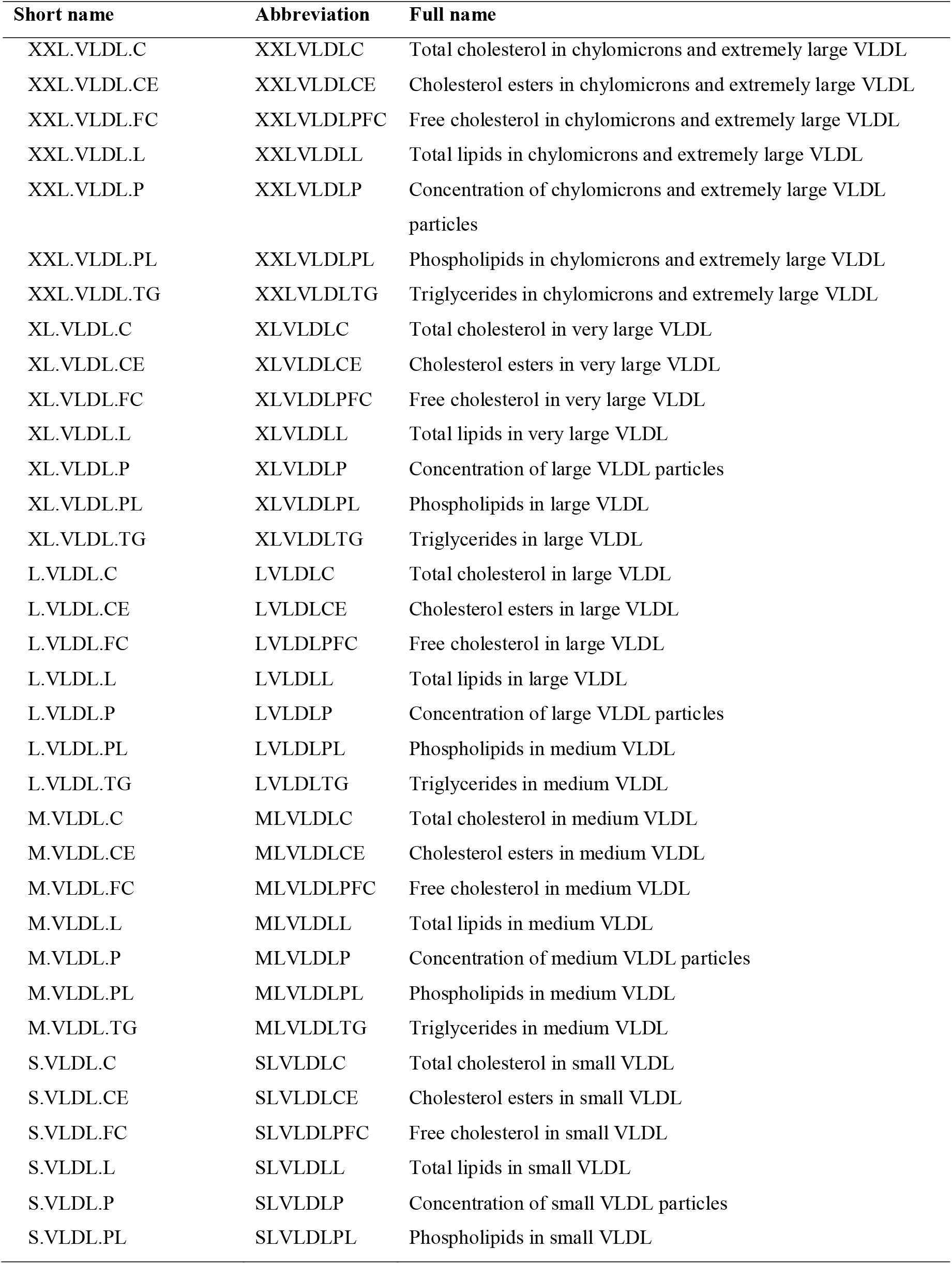
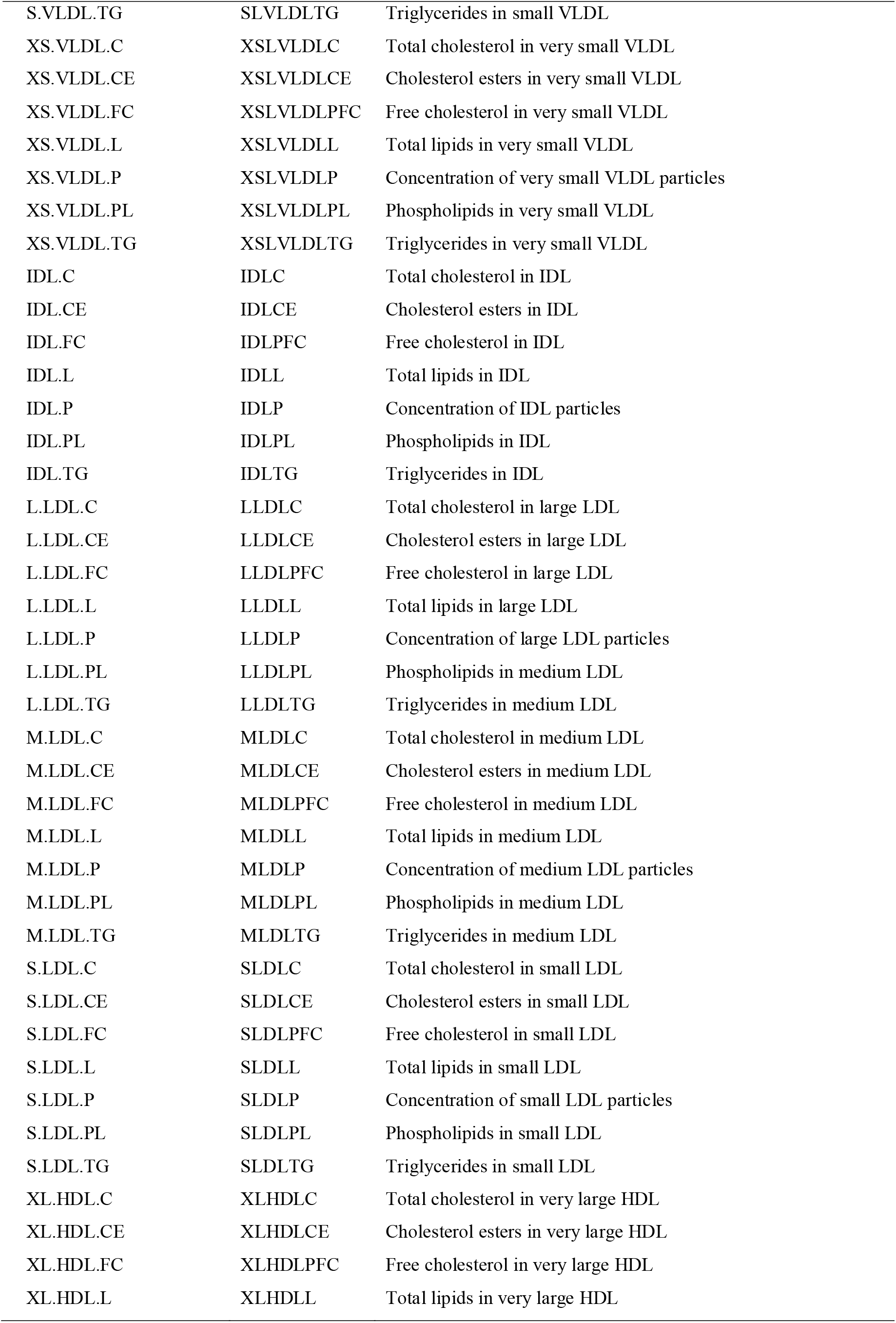
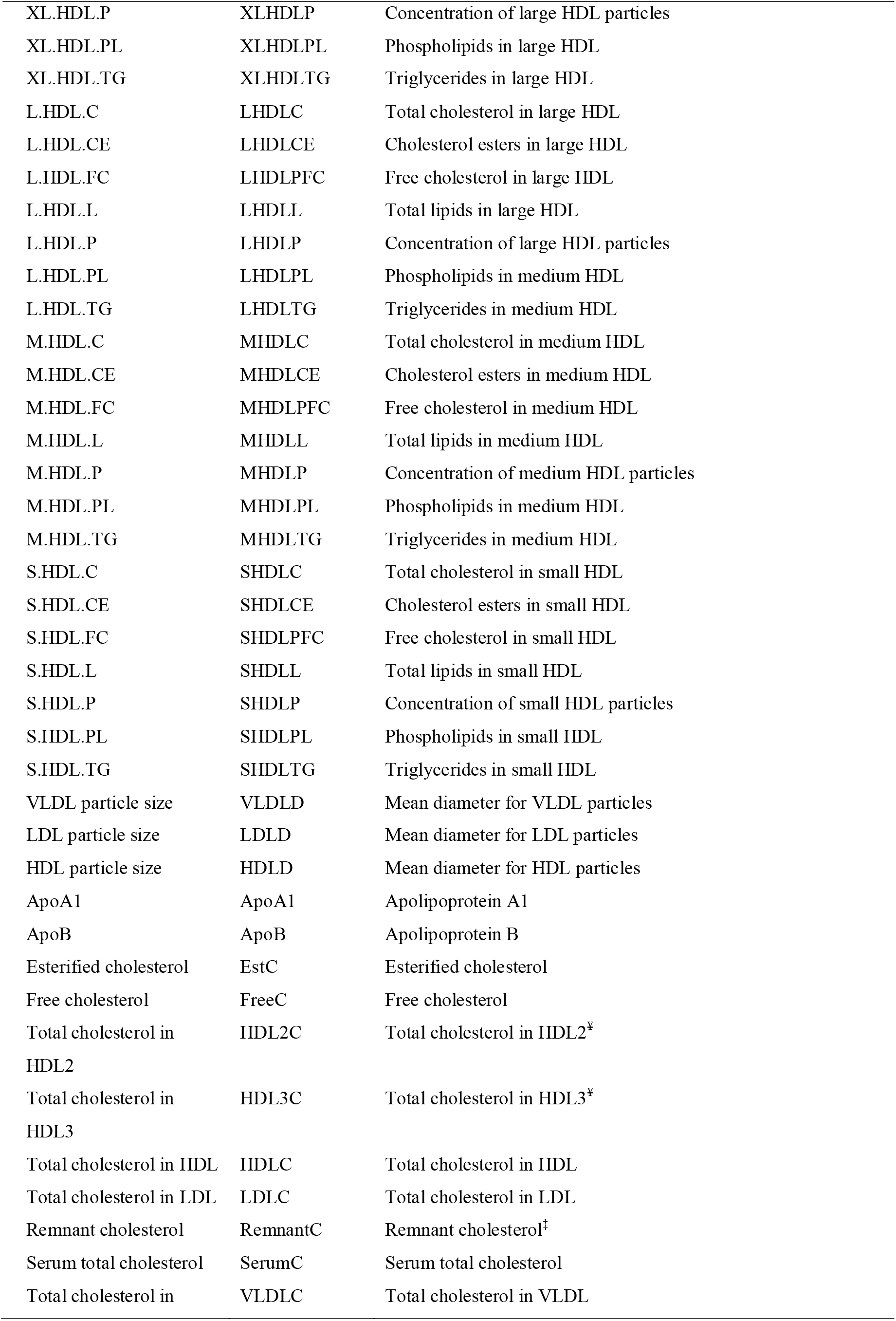
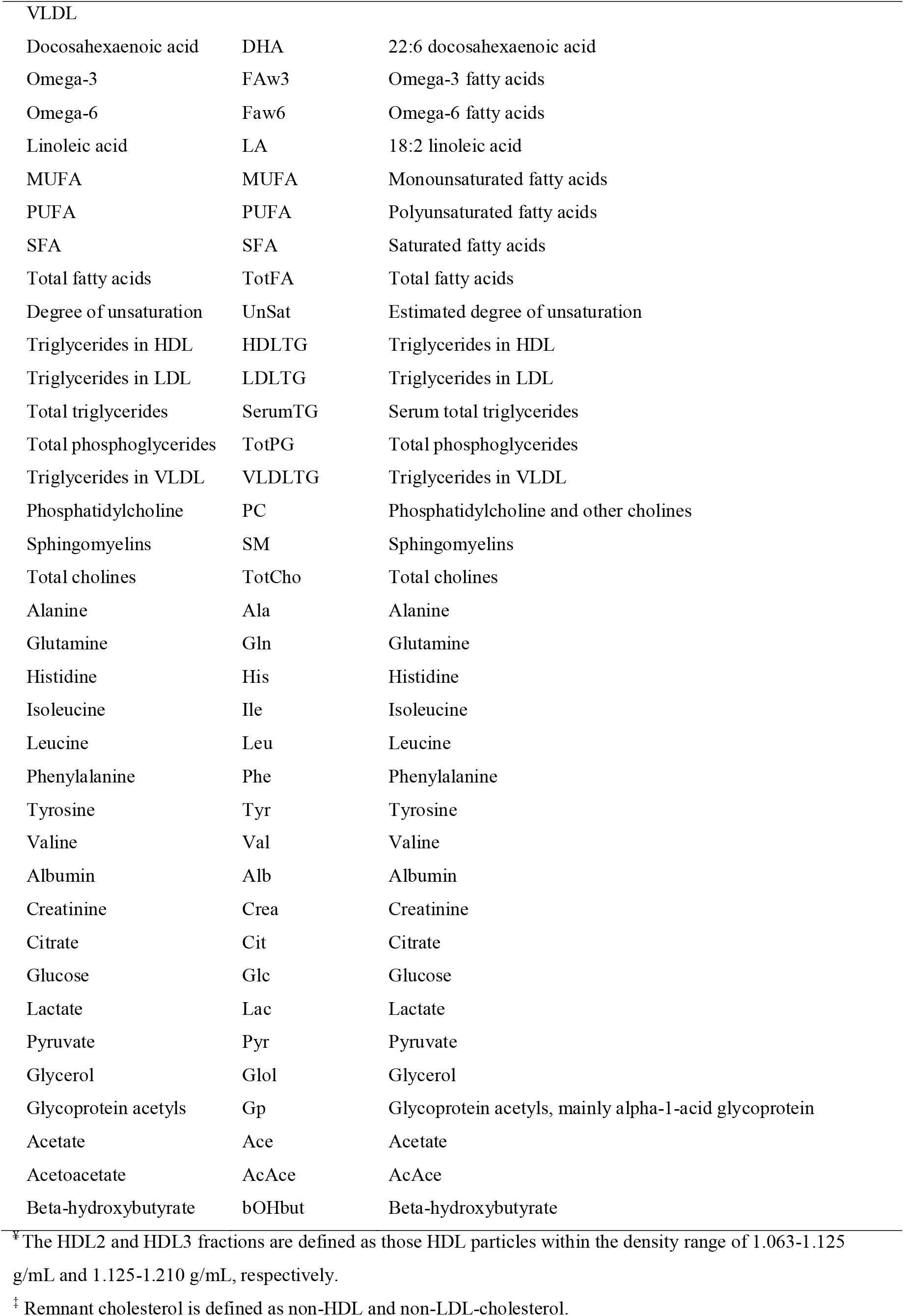
Description of 148 metabolic measures that were measured with a high-throughput proton nuclear magnetic resonance metabolomics platform (Nightingale Health Ltd., Helsinki, Finland).

